# Characterizing the Morphology of Costa Rican Stingless Bees to Parameterize the InVEST Crop Pollination Model

**DOI:** 10.1101/2022.10.07.511273

**Authors:** Christopher Sun, Rebecca Chaplin-Kramer

## Abstract

The InVEST Crop Pollination model operates on land use and land cover (LULC) characteristics, using available nesting sites and floral resources within a specified flight range to gauge the abundance and yield of bees species. In this study, we parameterize the InVEST Crop Pollination model to validate predictions of relative pollinator abundance in Costa Rica. Flight ranges of bee species are required as model inputs, yet are not readily available in literature compared to morphological attributes such as body length. To harness the availability of morphological data, we express the flight range of any given species as a function of its morphological attributes through a series of regressions, allowing for the estimation of flight ranges of species for which this metric is unknown. After proper parameterization, the model-predicted relative pollinator abundances of three species—*Tetragonisca angustula, Partamona orizabaensis, and Trigona corvina*—are compared against field data. A single proto-pollinator is then constructed as a representative species for analysis at a broader level, with model predictions validated against the total pollinator abundance across the entire spatial distribution represented by the field data. The model performs with a higher accuracy on the proto-pollinator compared to the individual species, revealing that there is surprisingly minimal added value from estimating individual flight ranges for each species. Rather, generalizing the biodiverse assortment of Costa Rican bees may yield better approximations for relative pollinator abundance.

## 1 Introduction

Land use and land cover changes are influencing the foraging behavior of pollinators. Crop pollinators are essential creatures because they provide crucial ecosystem services [1]. In Central American regions such as Costa Rica, changing proportions of forest cover and pastureland are altering the suitability of habitats for pollinators [2]. In turn, this impacts the quality of ecosystem services. Therefore, there is a need for the accurate LULC-based modeling of pollinator species to understand their pollinators’ response to habitat variability, predict future abundance trends, and properly inform political avenues of ecosystem repair to allow for the conservation of these species.

### 1.1 InVEST Crop Pollination model

The pollination model utilized in this study was the InVEST^1^ Crop Pollination Model, a part of the suite of InVEST models designed by the Natural Capital Project, which is headed by the Stanford Woods Institute for the Environment.

The InVEST Crop Pollination model operates on land use and land cover (LULC) characteristics, using available nesting sites and floral resources within a specified flight range to gauge the abundance and yield of bees species. As inputs, the model requires a guild table, a biophysical table, farm shapefiles, and an LULC raster for the region of interest. The biophysical table consists of a list of all habitat classifications and corresponding indices of cavity/ground nesting availability and floral resources availability. The guild table consists of a list of all pollinator species and four corresponding values: ground nesting suitability, cavity nesting suitability, flight range, and relative abundance. Using these inputs, the InVEST Crop Pollination model operates spatially over the region of interest and returns georeferenced rasters displaying gradients of pollinator abundance, pollinator supply, and wild/total pollinator yield.

The model’s user guide, documentation, and computational methodology can be found at [this link (click)](http://releases.naturalcapitalproject.org/invest-userguide/latest/croppollination.html).

### 1.2 Flight Range Dilemma

Pollinator flight range will be a subject of elaborate discussion in this article, as this metric was not widely available for all bee species. Hence, a necessary task before validation of the InVEST model was the deduction of the flight ranges of bee species for which data was unavailable. This task was treated as a supervised learning problem, explained in Section 2.1.

### 1.3 Objectives

The overarching objective of the research described in this article is to validate InVEST model predictions of pollinator abundance against field observations across four Costa Rican administrative divisions: San José, Heredia, Cartago, and Alajuela.

To accomplish this, open-source data is scraped from previously published studies through a systematic review, the results of which are used to form a data set of the morphological traits of different stingless bee species. Linear and polynomial regressions are employed to express flight range as a function of morphological variables.

These flight ranges allowed for the parameterization of the InVEST model to generate species-specific abundance predictions that were subsequently compared to field observations. We then apply various techniques and alter model inputs to optimize the performance of the InVEST model, maximizing adherence to ground truths. Ultimately, the construction of a proto-pollinator and the discovery of a relationship between pollinator abundance and elevation reveal the strengths and weaknesses of the model, pointing to future improvements in the crop pollination modeling framework.

## 2 Methodology: Flight Range Prediction

### 2.1 Systematic Review

The species chosen for the analysis were *Tetragonisca angustula, Trigona corvina, Partamona orizabaensis*, since the validation data contained the most observations for these species. Initially, we conducted a systematic review of the scientific literature pertaining to the flight range of these species. However, flight range data was only available for *T. angustula*. During the review, it was noted that morphological data on stingless bee species was significantly more abundant than flight range data. This observation motivated the hypothesis that the flight ranges of stingless bees could be predicted from morphological features. Hence, the objective of the literature review shifted towards identifying stingless bee species present in Costa Rica that had available data on flight range as well as morphology, and compiling these measurements to form a data set for analysis. Along with flight range, five morphological features were recorded: lowest observed body length in millimeters, highest observed body length in millimeters, body width (either intertegular span or thorax width) in millimeters, body mass in milligrams, and forewing length in millimeters. Flight ranges were recorded in meters.

Papers included in the review were all published in reputable journals. These literature were found by searching for key terms on the Google Scholar database. Experimental results and data measurements were extracted from these sources and compiled on a cumulative spreadsheet. Ultimately, we assembled a data set consisting of 22 species. Though all selected species contained measurements for flight range, not all species contained measurements for each of the five stated morphological variables; at this stage, there were still missing values in the data set.

### 2.2 Data Preprocessing

Body length and body width were the most abundant features, while body mass and forewing length contained more missing values. To impute these missing data points, the K-Nearest Neighbors model with a k-value of 3 was employed as an imputation mechanism. Missing values for a particular species were replaced by the weighted mean of the three nearest species in terms of Euclidean distance.

Following imputation, an exploratory analysis of the data revealed the presence of outliers, namely the unusually large documented flight ranges of Apis mellifera, Melipona beecheii, and Melipona subnitida. These outliers were removed using the IQR Rule and verified through Principal Component Analysis (PCA). PCA showed that the stated species formed a stray cluster in context of the data set, supporting the outlier identifications obtained through the IQR Rule.

Following KNN imputation and proper outlier removal, the data contained 19 records of the flight range and morphological characteristics of Costa Rican stingless bee species.

### 2.3 Morphological Analysis

The task at hand was concerned with establishing a generalized relationship between species’ morphology and flight range, allowing for the prediction of the flight range of Trigona corvina and Partamona orizabaensis, the original species of interest without documented flight ranges.

Single-variable linear regressions were implemented between each of the five morphological variables and flight range. The results of linear regression motivated a series of polynomial regressions for nonlinear modeling of the data, to achieve better fits and lower the magnitude of residuals. Finally, an ensemble methodology was employed to integrate the predictions of each individual regressor, dictating the final flight range prediction of a given species.

## 3 Results: Flight Range Prediction

Statistical analysis showed that body length and body width were strongly correlated with flight range, while body mass and forewing length were moderately correlated with flight range. The correlation between forewing length and flight range was negatively influenced by two data points with high leverage, corresponding to the species Melipona compressipes and Melipona quadrifasciata. Overall, forewing length constituted the weakest correlations with other morphological variables.

### 3.1 Regression Analysis

The linear relationship between body width and flight range achieved the highest degree of statistical significance, while that between forewing length and flight range achieved the lowest degree of statistical significance. Still, all linear relationships were significant at the *α* = 0.05 level. The equations of flight range as a linear function of morphological attributes are listed below.

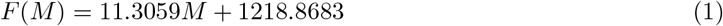

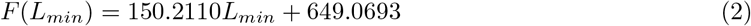

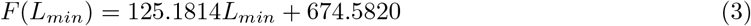

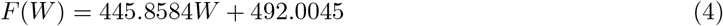

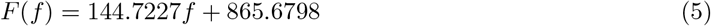

Upon closer examination of the distribution of the two-variable data as well as the residual plots of the linear models, it was determined that the data conformed to a nonlinear relationship, slightly concave downwards relative to the positive axes. Theoretically, this observation was justified, as described by Grula et al. (2021), who concluded that the allometry inherent to bee morphology creates an inversely proportional relationship between body size and wing length [3]. As smaller bees are less equipped for extensive flight, these phenomena may create the dip near the extreme values of the distribution. The polynomial relationship was observed for all features except forewing length. Hence, second-order polynomial regression was conducted on the same scatterplots with the exception of forewing length, the results of which are displayed in Figure 3.

**Figure 1:**
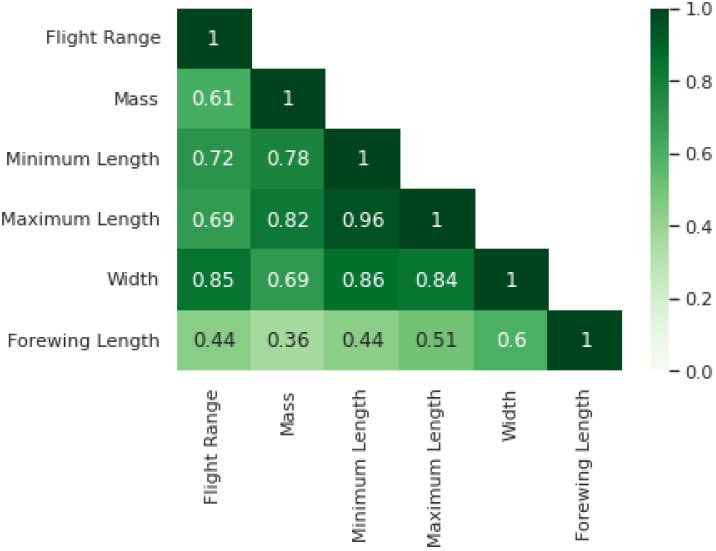
Correlation Matrix

**Figure 2:**
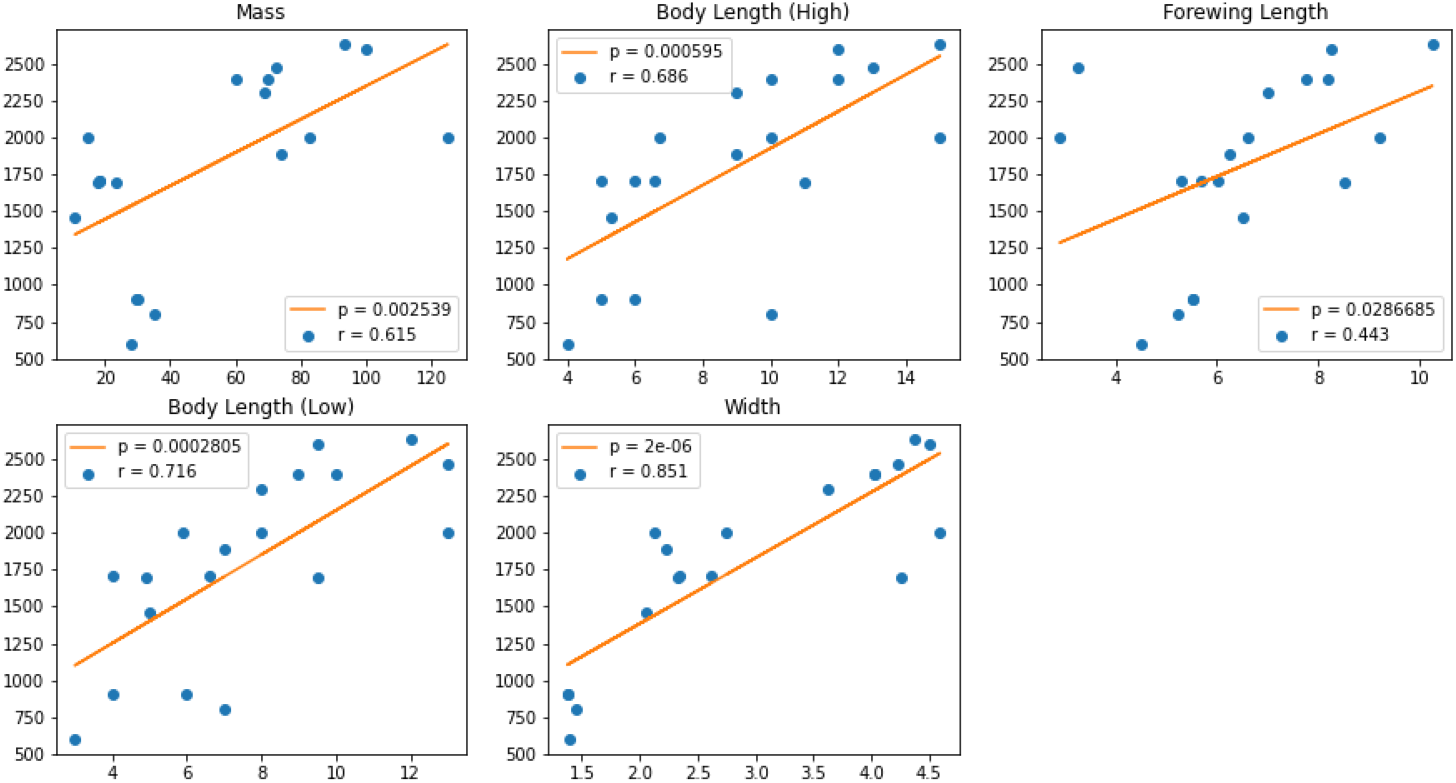
Linear Models Relating Morphology to Flight Range

**Figure 3:**
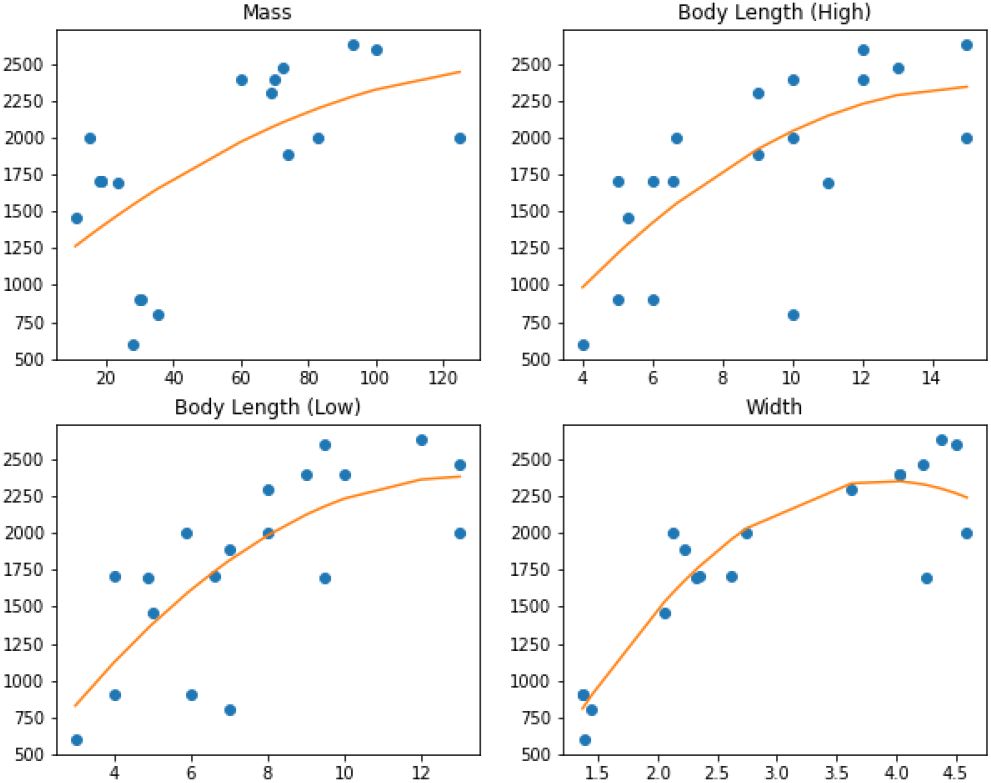
Nonlinear Second Order Polynomial Models Relating Morphology to Flight Range

According to the polynomial relationship between flight range and morphological attributes, once a given species reaches a threshold measurement, be it mass, length, or width, the corresponding flight range of the species starts to plateau, increasing at a decreasing rate. In fact, flight range is even projected to decrease at an increasing rate after the body width of a given species reaches 3.905 millimeters. The equations of flight range as a quadratic function of morphological attributes are listed below.

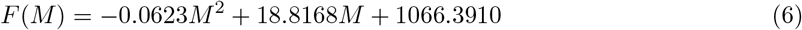

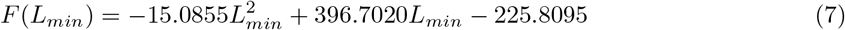

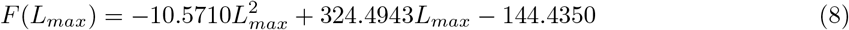

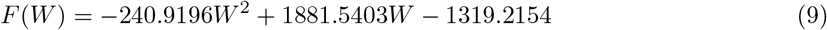

### 3.2 Comparative Correlation Analysis

Across all morphological features studied, a quadratic fit of the data achieved a lower mean absolute error than a linear fit. For all features except body mass, a quadratic fit of the data resulted in a lower mean absolute percentage error than a linear fit. However, these differences in error were not statistically significant, as seen by the large *P_MAE_* values in Table 1.^2^ Hence, to predict the flight ranges of the species of interest, T. corvina and P. orizabaensis, an ensemble approach was used, weighing the predictions of each individual regressor equally to output a final prediction of flight range. This process is represented as follows:

**Table 1:** Caption

| | Linear | | Polynomial | | $P_{MAE}$ |
| --- | --- | --- | --- | --- | --- |
|  | MAE (m) | MAPE (%) | MAE (m) | MAPE (%) |  |
| Mass | 423.6 | 0.248 | 422.9 | 0.252 | 0.4699 |
| Length (Low) | 369.6 | 0.221 | 322.2 | 0.196 | 0.3884 |
| Length (High) | 371.9 | 0.222 | 350.2 | 0.210 | 0.4648 |
| Width | 261.7 | 0.163 | 179.1 | 0.107 | 0.2280 |
| Forewing Length | 446.2 | 0.270 | - | - | - |

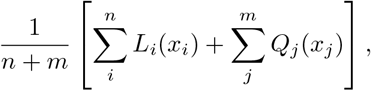

with *i* and *j* being the morphological attribute subject to the corresponding linear model *L*(*x*) and the corresponding quadratic model *Q*(*x*), respectively.

Multivariable regressions were also used to model the data with superior results, but due to the absence of data on some morphological features for the target species, these techniques were omitted as impractical solutions to the flight range dilemma.

### 3.3 Unknown Flight Range Predictions

Using this outlined framework, T. corvina is predicted to have a flight range of 1,399.7 meters, and P. orizabaensis is predicted to have a flight range of 1,534.4 meters.

## 4 Pollinator Abundance Validation

### 4.1 Species-Specific Validation

The InVEST Crop Pollination model was run on the three species of interest. Let a sequence of relative individual pollinator abundances be defined as

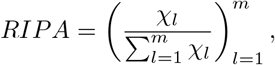

where *χ_l_* is the pollinator count at the *l^th^* location for a total of *m* locations. For each species, the model’s prediction of relative pollinator abundance (RIPA) was compared to its true RIPA at every location containing observations of that species in the field data. Shown in Figure 4 are plots for each species regressing these true RIPA values against predicted values. Zero-valued predictions and statistical outliers for both true and predicted values were removed from the analysis, as they deflated correlations and rendered them unrepresentative of the model’s performance.

**Figure 4:**
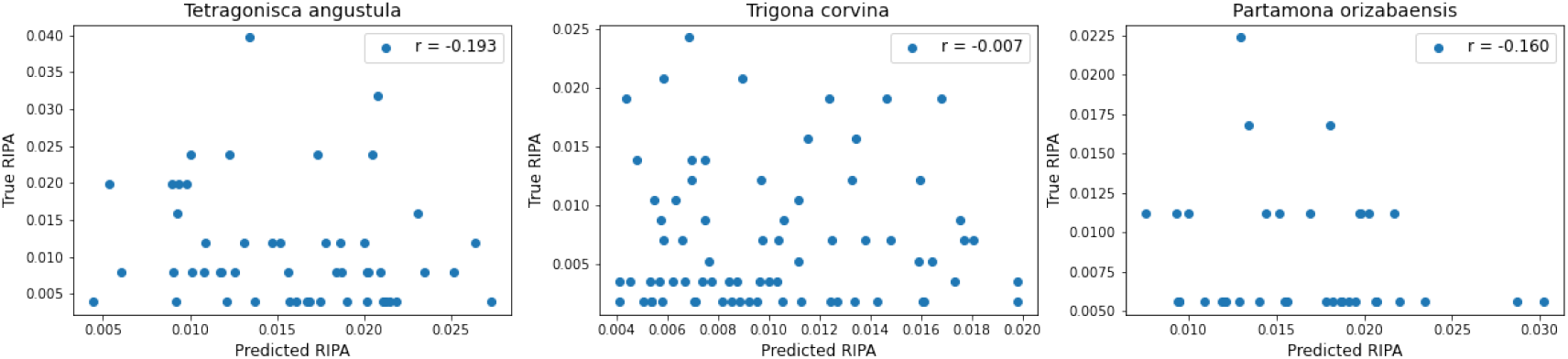
Species-Specific Abundance Validation Results

These species-specific results show that the InVEST model performed poorly, unable to correctly project RIPA values that align with field data. Given these suboptimal results, we were faced with two possible explanations.

1. The field data, when taken in the narrow context of a single species, presented too much variability where the validation RIPA was not trustworthy, hence the discrepancy between true and predicted values even if the model had integrity.
2. The model itself was inaccurate; even at broader multi-species levels, a suboptimal correlation between true and predicted RIPA would occur.

To test these conjectures, we shifted from species-specific validation toward validation on the basis of broader groups of species. An explanation of our approach requires us to define the term “proto-pollinator.”

### 4.2 Proto-Pollinator Total Abundance Validation

Let “proto-pollinator” be defined as a hypothetical species that is morphologically and ecologically representative of all the species in an area. As opposed to species-specific analyses, the abundance predictions of a proto-pollinator are symbolic of *total pollinator abundance* (RTPA). Because the proto-pollinator is representative of species diversity, the fact that there are different species existing at each location can be neglected, and proto-pollinator abundance at any location can be equated to the RTPA at that location.

We wanted to construct a proto-pollinator and feed its attributes to the InVEST model to compare model outputs to total pollinator abundance values. A subsequent comparison of species-specific results and proto-pollinator results would allow us to gauge the added value of conducting in-depth analyses for each species and finding their individual flight ranges. Most importantly, an establishment of this added value as insignificant would allow for the conclusion that in-depth species-specific analyses are not worth the effort and do not yield results considerably better than broader analyses.

The proto-pollinator we constructed was assigned a 1,600-meter flight range and nesting suitability in both cavity and ground habitats. Resulting outputs of proto-pollinator abundance were considered as measurements of RTPA relative to each location.

Let a sequence of RTPA measurements relative to *m* locations be defined as:

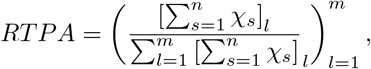

where *χ_s_* is the pollinator count for species *s*, *n* is the number of species at each of the *m* locations, and *l* is an individual location. *RTPA* contained the ground truth values compared to model outputs. The plots in Figure 5 demonstrate a weak to moderate correlation between predicted and true RTPA. Correlation coefficients of proto-pollinator validation are considerably higher than the those of single-species validations. The improvement that comes with proto-pollinator modeling relative to single-species modeling demonstrates that the InVEST Crop Pollination model may perform more accurately with less (but broadly-grounded) information than with more (but sparsely-grounded) information about pollinators.

**Figure 5:**
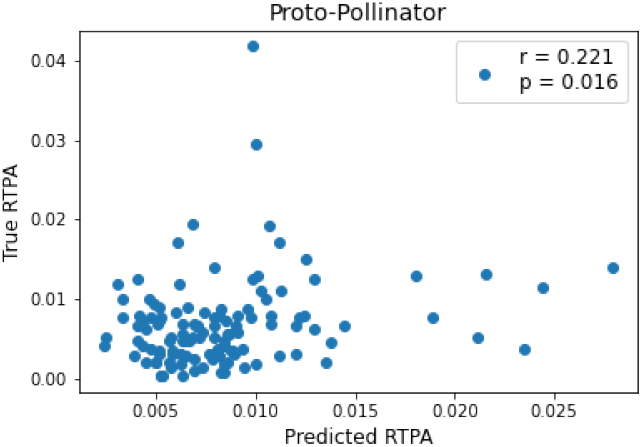
Proto-Pollinator Abundance Validation Results

**Figure 6:**
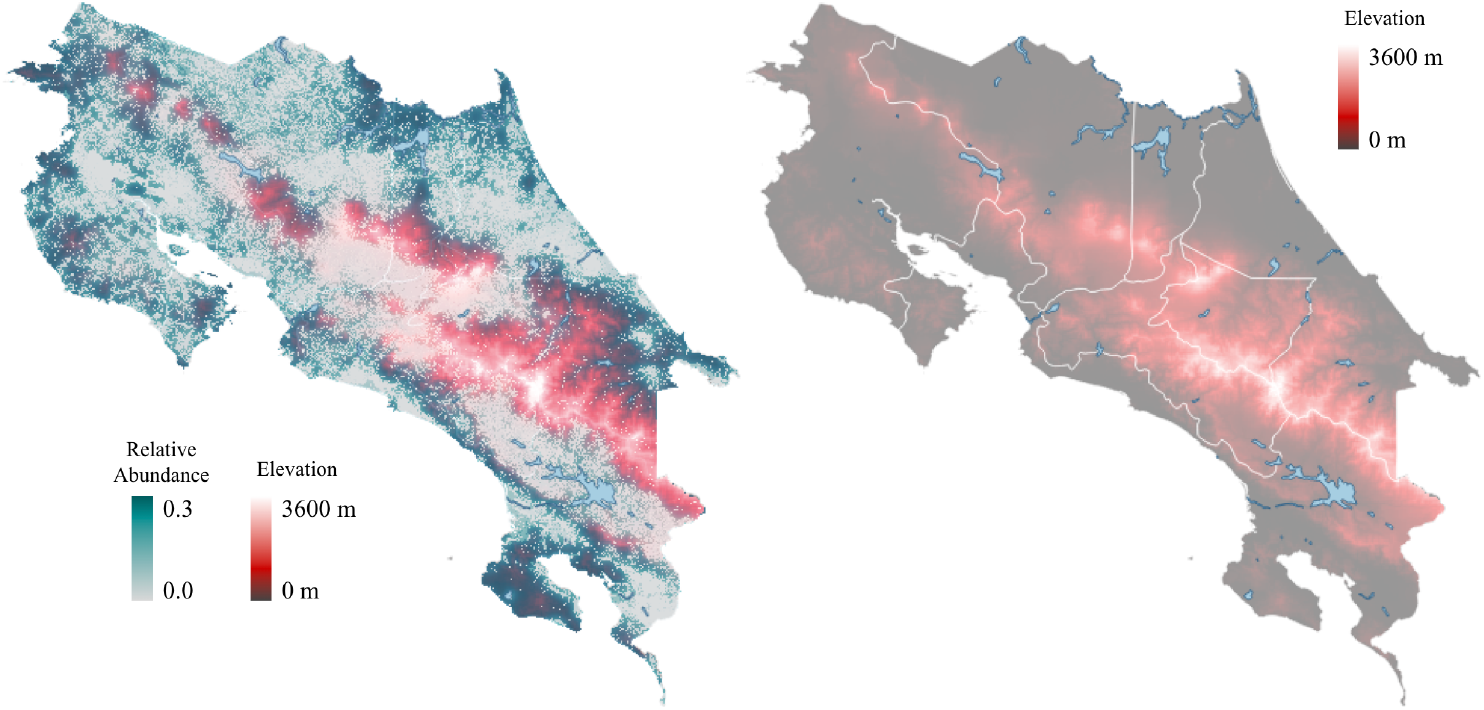
Relationship Between Elevation and Abundance Predictions

**Figure 7:**
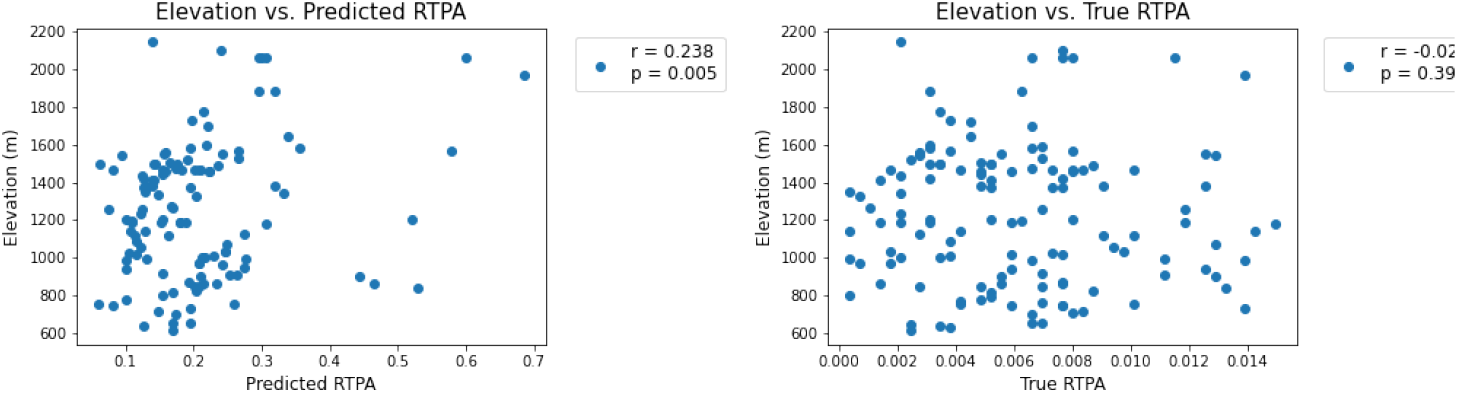
Discrepancy in Relationship Between Elevation and Abundance for Predictions and Ground Truths

**Figure 8:**
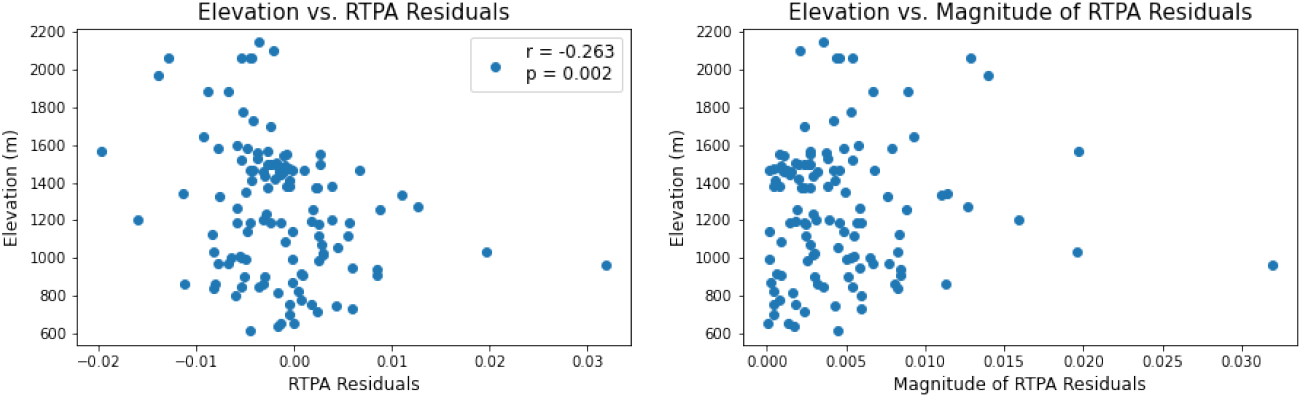
Relationship Between Elevation and Abundance Residuals

Referring back to the explanation postulated in Section 4.1, we have demonstrated that the first explanation holds true in the context of Costa Rican pollinators. Modeling on larger scales and in other geographic regions is needed to gauge the degree to which this explanation holds true in different contexts.

### 4.3 Confounding Impacts of Elevation

While not evident from raw outputs, the InVEST Crop Pollination model associates relative pollinator abundance levels throughout Costa Rica with elevation. Namely, regions with higher altitudes contain a higher relative abundance compared to low altitude counterparts. This observation is surprising, since the InVEST framework does not explicitly account for elevation.

The topography of Costa Rica is relatively simple; regions along the coastlines have low elevations while mountain ranges extending NW-SE reach over 3,000 meters in elevation, providing high-altitude habitats for pollinators. According to the model, pollinators tend to prefer the northeast face of the Talamanca Mountains (Cordillera de Talamanca), with the habitat range extending into the Panama border and to the Caribbean coast. Meanwhile, in valley regions, such as southwest of the Talamanca Mountains and between the Talamanca and Central ranges, abundance levels are low. This pattern is also observed in the Tilarán Mountains (Cordillera de Tilarán), with the immediate region surrounding the mountain range having low abundance levels but the range itself having higher abundance levels.

We proceeded to explore these observations numerically by acquiring the elevation in locations represented by abundance predictions. Regressing these two variables revealed a moderate positive correlation. However, there is no relationship between ground truth pollinator abundance and elevation. Hence, we suspect the model is operating on a parameter confounded with elevation.

Our hypothesis for the fact that there is a positive correlation between elevation and predicted pollinator abundance is that elevation is confounded with availability of nesting habitats and quality of climate. Namely, the proportion of land cover that is forest increases at higher-altitude regions in Costa Rica (citation needed). This would increase the availability of nesting, which is accounted for by the model given predictions of higher relative pollinator abundance in these higher-elevation areas. However, the model does not account for the tradeoff that comes with higher elevations: a decline in climate suitability (citation needed). If this was the case, there would be no relationship between elevation and abundance in reality, supporting field observations.

#### 4.3.1 Proto-Pollinator Residuals at Varying Elevations

To test our hypothesis that the model accounted for only one side of the climate-habitat tradeoff, we regressed elevation against the residuals of proto-pollinator abundance. The resulting plot showed a moderate negative correlation.

At higher elevations (1,600 meters and above), residuals were all negative, representing an overestimate of pollinator abundance. At medium and lower elevations, residuals were a mix of positive and negative, with more instances of large positive residuals at lower elevations (around 1,000 meters). These findings support our hypothesis, as an overestimation of floral resources and nesting suitability would lead to overestimations of pollinator abundance.

Considering these analyses, the confounding nature of elevation reveals a possible shortcoming of the InVEST framework and crop-based models in general that do not account for climate suitability in varying environments.

### 4.4 When To Take the Average?

This section will explore the fundamental question of whether pre-result or post-result generalization of species-specific diversity leads to better modeling performance in the context of Costa Rican pollinators. Given the superior validation results of proto-pollinator modeling compared to species-specific modeling, it becomes natural to question whether the aggregate abundance of a set of species achieves better correlations than individual abundance predictions. We will define “aggregate abundance” as the sum of abundance across a set of species. Aggregate abundance is not equivalent to RTPA, as the latter is the total observation counts of pollinators without regard for species. Particularly, aggregate abundance is a fraction of RTPA and is defined this way for the purposes of testing whether results of summing certain representative species can simulate RTPA even when those species account for only a fraction of species known to exist. In other words, if aggregate abundance predictions achieve higher correlations than species-specific predictions, it becomes necessary to also determine whether aggregate abundance predictions conform to original RTPA ground truths. Fulfillment of both conditions would provide evidence for the claim that generalization leads to better validations.

In the field data, there were 26 locations containing observations of all three species of interest: *T. angustula, T. corvina, P. orizabaensis*. At each of these locations, predicted and true RIPA for the individual species were summed and compared. Surprisingly, a moderate correlation was discovered between predicted and true values of aggregate abundance. Recall that species-specific correlations were suboptimal, with some species even having negative correlation coefficients. The considerable increase in accuracy associated with a species-wise totaling of RIPA provides insight into the added value of species generalization.

We have already established that RTPA validation results are superior those of RIPA. The task at hand is to establish that RAPA validation results are also superior than those of RIPA, but inferior to those of RTPA. Figure 9 accomplishes both aspects of this task.

**Figure 9:**
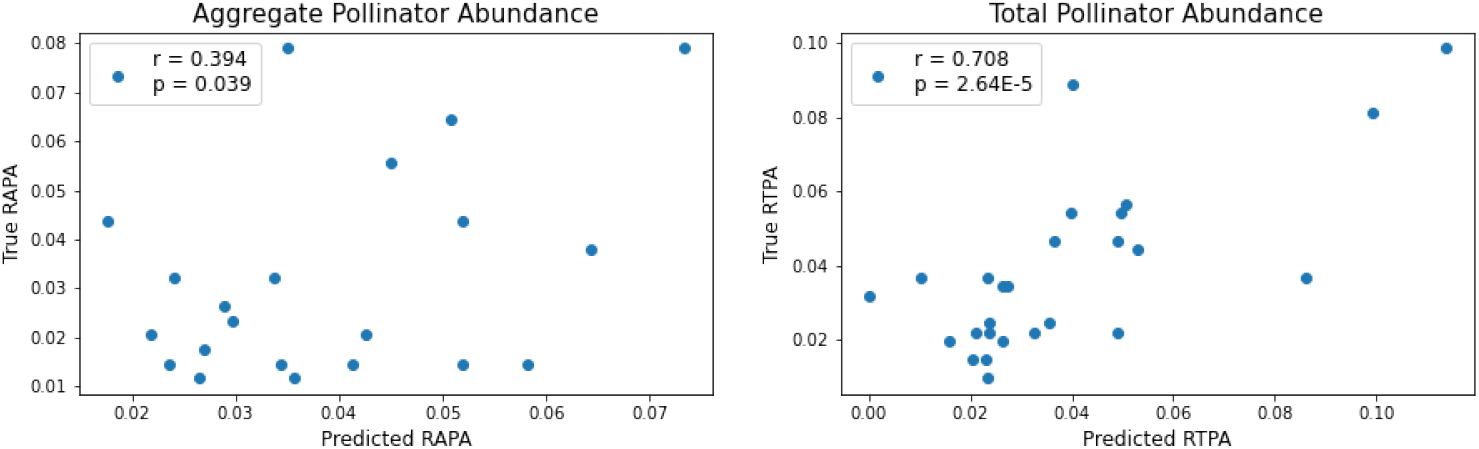
Aggregate Abundance Validation Results

The following series of expressions symbolically conveys the overall motivation for our modeling methodology. In all expressions, let the fraction operator be interpreted as an element-wise comparison between the numerator and the denominator for an ideal one-to-one correlation. Let *x_n_* represent the relative abundance for the *n^th^* species relative to *m* locations.

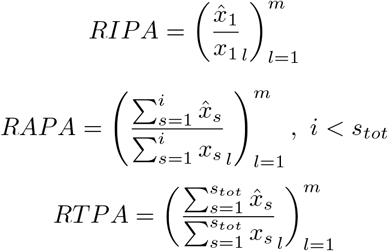

Furthermore, assuming

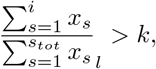

for a certain threshold value *k* that would allow for the assumption of representativeness, is it possible to show that

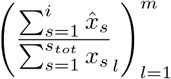

is comparable to or even outperforms RTPA?

## 5 Conclusions

In ecological modeling, researchers often find it hard to characterize parameters either due to scarcity of necessary metrics or unavailability of literature on the topic of interest. In the context of the InVEST Crop Pollination Model, biophysical and guild tables present this challenge, as information regarding the flight range and nesting suitability of pollinators, as well as the nesting and floral resources availability, of varying land cover types, are not readily available. Specifically, acquiring flight range values is time-consuming, hence the need to develop a streamlined way to infer flight range from morphological characteristics. In this study, we have conducted systematic reviews to search for morphological data exhaustively, allowing us to devise linear and polynomial relationships to predict the flight range of any given species, as long as simple anatomical information about the species is known.

Establishing these relationships allowed us to conduct single-species validations on model-predicted relative individual pollinator abundance (RIPA) values for three chosen species: *T. angustula, T. corvina, and P. orizabaensis*. However, suboptimal results and inadequate correlations prompted us to rethink the task of abundance modeling. We subsequently turned to a series of generalizations, constructing a proto-pollinator representative of all species in the region, and feeding its traits into the model to generate relative total pollinator abundance (RTPA) predictions. Results showed substantial improvements, with a significantly higher degree of adherence between predicted and true values. Creating a proto-pollinator also prompted the discovery of elevation’s confounding nature as a variable that is not considered by the model yet is associated with model predictions.

We further explore the notion of generalization using species aggregation, in which we sum both the predicted and true RIPA values of the three chosen species to calculate relative aggregate pollinator abundance (RAPA) values. Surprisingly, correlations between predicted and values improve significantly as a product of these summations.

In a way, the results of this study invalidate the need to model species individually and determine parameters such as flight range. The tedious workflows that come with species-specific modeling may very well result in suboptimal validation results. On the other hand, generalizing traits of species in the form of aggregate abudance or proto-pollinator abundance may be the more time-efficient and accurate choice.

## Footnotes

1 Integrated Valuation of Ecosystem Services and Tradeoffs

2 P-values were obtained through resampling procedures, as the distribution of error across all species was not normal.

